# Laminar fMRI in the locked-in stage of amyotrophic lateral sclerosis shows preserved activity in layer Vb of primary motor cortex

**DOI:** 10.1101/2024.04.26.590857

**Authors:** Lasse Knudsen, Bo Jespersen, Mia B. Heintzelmann, Peng Zhang, Yan Yang, Torben E. Lund, Jakob U. Blicher

## Abstract

Amyotrophic lateral sclerosis (ALS) affects the cerebral cortex layer-dependently, most notably by the foremost targeting of upper motor neurons (UMNs) sited in layer Vb. Previous studies have shown a retained ability of paralysed patients to activate residual cortical motor networks, even in late-stage ALS. However, it is currently unknown whether such activation reflects a retained capacity to process sensorimotor inputs or if it is a result of actual motor output. Given the distinct function of individual cortical layers, layer-specific functional measurements may provide insight to this question. In this study, using submillimetre resolution laminar fMRI, we assessed the layer-dependent activation associated with attempted (motor) and passive (somatosensory) movements in a locked-in stage ALS patient. We found robust activation in both superficial and deep layers of primary motor cortex. The peak activation in deep layers was localised to layer Vb. These findings demonstrate preserved activity in deep output layers of M1, possibly reflecting a retained ability to engage residual UMNs despite years of paralysis. Our study underscores the capacity of laminar fMRI to discern subtle cortical activity and elucidates a promising pathway for probing in vivo human ALS pathology with unprecedented resolution.

## Introduction

Amyotrophic lateral sclerosis (ALS) is a fatal neurodegenerative disease, characterised by progressive loss of both spinal lower motor neurons (LMNs) and cortical upper motor neurons (UMNs)^1^. Respiratory failure is the primary cause of mortality, but some countries optionally provide invasive tracheostomy, which can prolong life considerably. Patients in this condition gradually progress to the locked-in stage, characterised by full paralysis and limited communication through eye movements or assistive devices. ALS affects the motor cortex layer-dependently^2^. That is, neurons in the deep output layers (hereunder UMNs in layer Vb) are believed to be affected first, from which degeneration progressively spreads towards superficial layers associated with integration of sensorimotor inputs (Fig. 1A). There is evidence to suggest that patients retain the capability to intentionally activate spared cortical motor networks even in the locked-in stage^3,4^. However, deciphering the layer-specific cortical contribution to such activity has been unattainable due to a lack of methodologies with adequate spatial resolution. The functional characteristics (e.g. input versus output activity) hence cannot be inferred, and it remains unknown whether late-stage ALS patients retain functionally capable neurons in the deep output layers. To this end, recent advances in functional magnetic resonance imaging (fMRI) have enabled acquisitions with submillimetre resolution, allowing for functional recordings from individual divisions of the cortical layers (laminar fMRI)^5–7^. The utility of this technique in assessing layer-dependent hypotheses in clinical populations was recently demonstrated in dystonia patients^8^. Considering the layer-specific neuropathology presented by ALS, McColgan and colleagues have advocated for its application in this condition^2^. Such data, however, is yet to be published.

**Figure 1.**
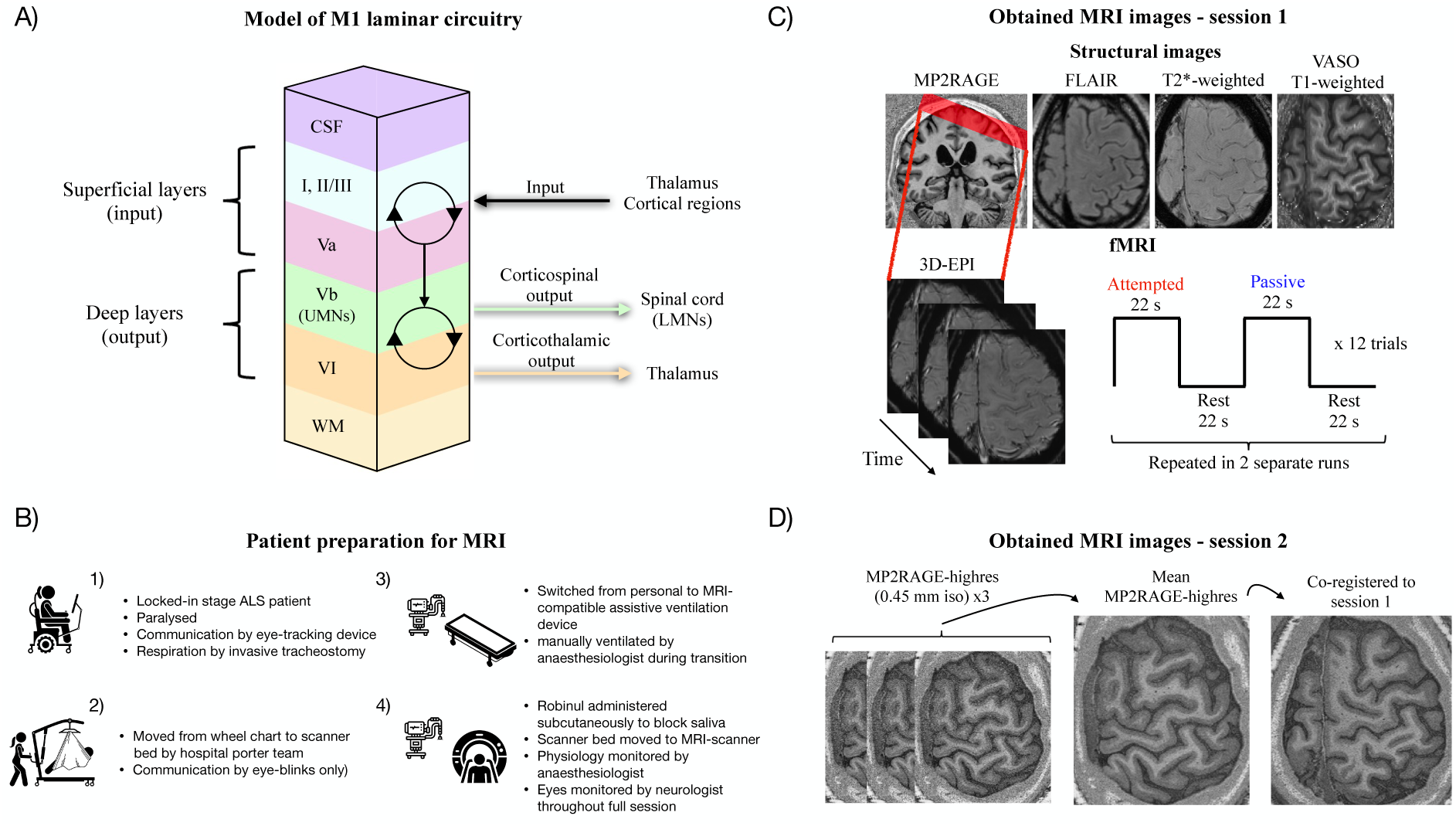
Overview of M1 laminar circuitry and the experimental setup used to scan the patient. (**A**) Laminar input/output organisation based on animal literature (rodents and non-human primates)*^9–15^*. The majority of sensorimotor input is integrated in superficial layers of M1 whereas corticofugal output emerge from the deep layers. Input information is believed to be pre-processed via recurrent activation in superficial layers before being relayed to deep layers to inform motor output (rings and arrows)*^16,17^*. Recent laminar fMRI studies in healthy subjects support a similar model in humans*^6,18–21^*. UMNs (layer Vb) and LMNs (spinal cord) are the cardinal cells affected by ALS (**B**) Overview of patient characteristics (details provided in Materials and Methods), and illustration of the procedures to overcome challenges of scanning a locked-in stage ALS patient. (**C**) Several structural images were obtained with different purposes. MP2RAGE was used to place the imaging slab for functional acquisitions, for grey matter segmentation and for co-registration with high resolution MP2RAGE images from session 2 (see (D)). The FLAIR and T2*-weighted structural images were evaluated for markers of neurodegeneration. The VASO T1-weighted image facilitated high-quality registration of MP2RAGE to EPI-space. BOLD-response measurements were obtained using 3D gradient-echo EPI (0.82 mm isotropic resolution). The functional paradigm was organised in blocks of attempted and passive finger movements alternating with rest. (**D**) The patient was scanned in a second session where MP2RAGE images with 0.45 mm isotropic resolution (MP2RAGE-highres) were obtained for approximate assignment of cortical layers with respect to the fMRI activation peaks obtained in session 1.

In this study, laminar fMRI was employed to evaluate mesoscale activation in the primary motor cortex (M1) of a patient in the locked-in stage of ALS (Fig. 1B-D). First, we evaluated functional responses to attempted (motor) and passive (somatosensory) finger movements in the hand area of M1. In subsequent laminar analyses, we aimed to demonstrate whether: 1) observed M1 activation during attempted movements mainly involved superficial layers as evidence of input-dominated processing due to depleted neurons in the deep layers, or 2) whether the deep layers were concomitantly engaged as evidence for preserved activity in the output layers of M1, possibly driven by residual functional UMNs.

## Results

We recruited a patient in the locked-in stage diagnosed with ALS six years prior (Materials and Methods). The patient was paralysed, reliant on artificial ventilation (invasive tracheostomy) and communication was only possible through an assistive eye-tracking device (Fig. 1B). In the main session, structural images for clinical characterisation were obtained in addition to the laminar fMRI acquisitions (Fig. 1C). The patient was subsequently invited for a second imaging session with the purpose of assigning cortical layers to functional activation peaks observed in session 1 (Fig. 1D). A healthy control was additionally recruited to supplement the main findings. This subject (HC) was scanned in a single session with an identical protocol as session 1 for the patient.

### Structural imaging-based characterisation of cortical pathology

The relative extent of cerebral and spinal involvement is highly heterogonous across ALS patients^22^. In some cases, severe cortical atrophy may follow from the aggressive neurodegeneration (see e.g.^23^), which would potentially render laminar analysis infeasible. Therefore, as a first step, the structural intactness of the patient’s cortex was assessed, which seemed largely intact with no clear signs of cortical atrophy based on MP2RAGE (Fig. 2A). Aside from atrophy, the patient had clear signs of cortical pathology, however, as evident from 1) clinical examinations in the early stages of the disease demonstrated extensive signs of UMN pathology (which were absent at the time of this study due to masking by progressed LMN depletion, see Materials and Methods); 2) high-resolution T2*-weighted images clearly showed the motor band sign (Fig. 2B), a well-known marker of UMN involvement^24^. The fluid-attenuated inversion recovery (FLAIR)-image did not show clear T2-weighted corticospinal tract hyperintensities (Fig. 2C), which has been proposed as a marker of axonal degeneration^24^. Its absence might thus be indicative of less affected corticospinal myelin, or could simply reflect a lack of sensitivity for this marker^24^.

**Figure 2.**
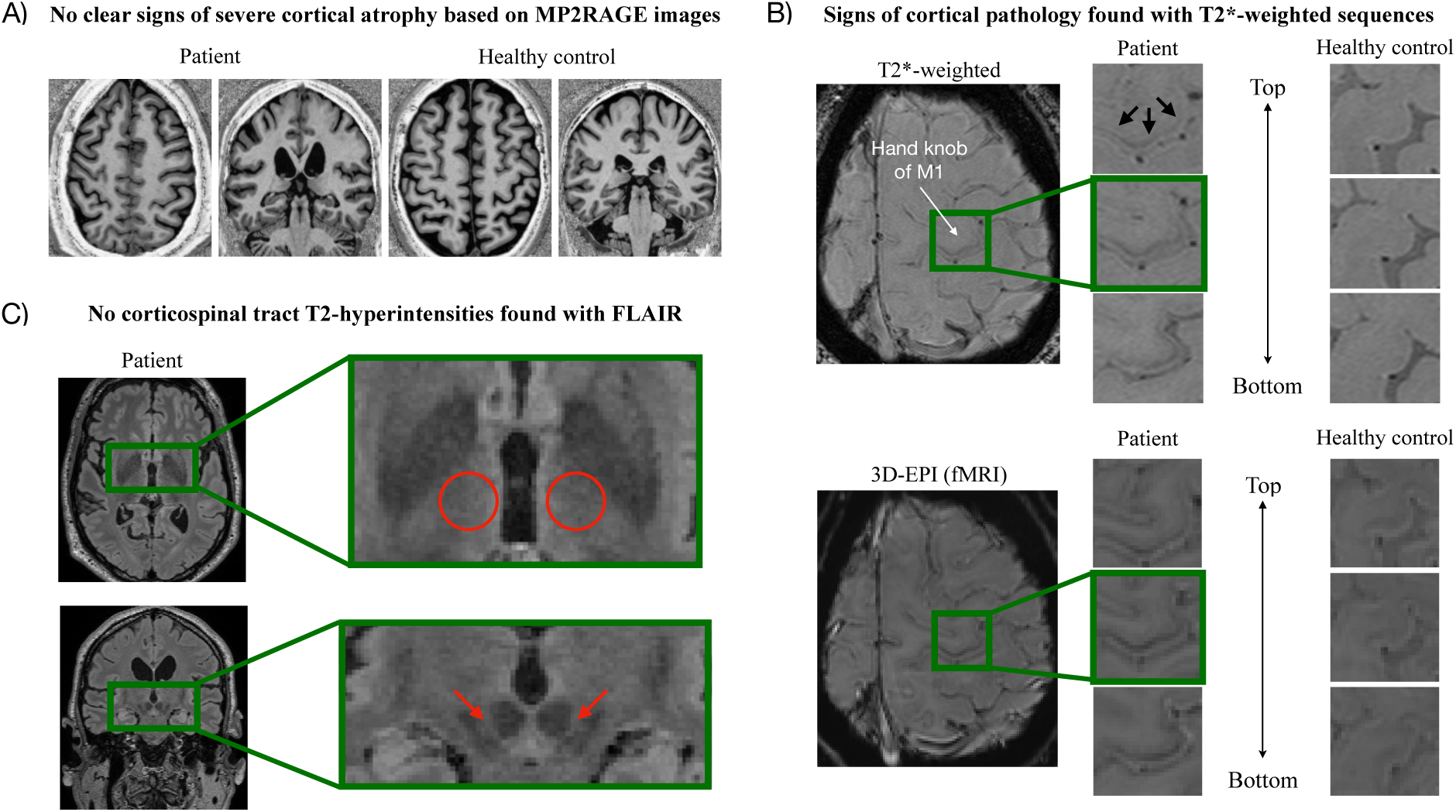
Structural imaging assessment of UMN involvement and atrophy. (**A**) Example axial and coronal MP2RAGE slices in the patient and healthy control. Considering the advanced stage of the disease, the patient’s brain appears remarkably similar to that of the healthy control. (**B**) The structural T2*-weighted image clearly demonstrated the motor band sign, i.e., a clear hypointense stripe (highlighted by arrows) suggestive of cortical pathology. Zoomed insets show this across three example axial slices at different heights in the precentral bank of the central sulcus. It was similarly visible in the mean 3D-EPI image of the patient (also T2*-weighted), and not present in the healthy control. (**C**) The FLAIR image revealed no clear T2-hyperintensities in the corticospinal tracts (insets highlight the expected bilateral positions with rings and arrows in example axial and coronal slices, respectively). Such hyperintensities have previously been suggested as a marker of cortical tract degeneration but may be insensitive*^24^*.

### M1 responses to attempted and passive finger movements

If the cortical motor networks of the patient were still functional, M1 should expectedly respond during attempted movements as seen for example in paralysed spinal cord injury patients^25,26^. Conversely, the primary somatosensory cortex (S1) is expected to not or only weakly activate given the lack of somatosensory involvement during non-overt movement. Since somatosensation is relatively unaffected in ALS^1^, both M1 and S1 were expected to activate during passive movements as in healthy subjects^21,27,28^. Blood oxygen level-dependent (BOLD) fMRI at 0.82 mm isotropic resolution was employed to evaluate cortical activation during these tasks (Fig. 1C). The resulting activation maps and region of interest (ROI) quantifications are shown in Fig. 3A. Attempted movements resulted in robust BOLD responses confined mainly to M1 (the mean difference in percent signal change compared with control regions was 0.59 percentage points (pp.) for M1 (CI95[0.55;0.63]) and 0.08 pp. for S1 (CI95[0.04;0.13])) (Table S1). Passive movements robustly activated both regions (Fig. 3A, Table S1). For HC, active and passive movements elicited significant activation of both M1 and S1 compared with control regions (Fig. S1, Table S1) in line with overt movement being present in both conditions.

**Figure 3.**
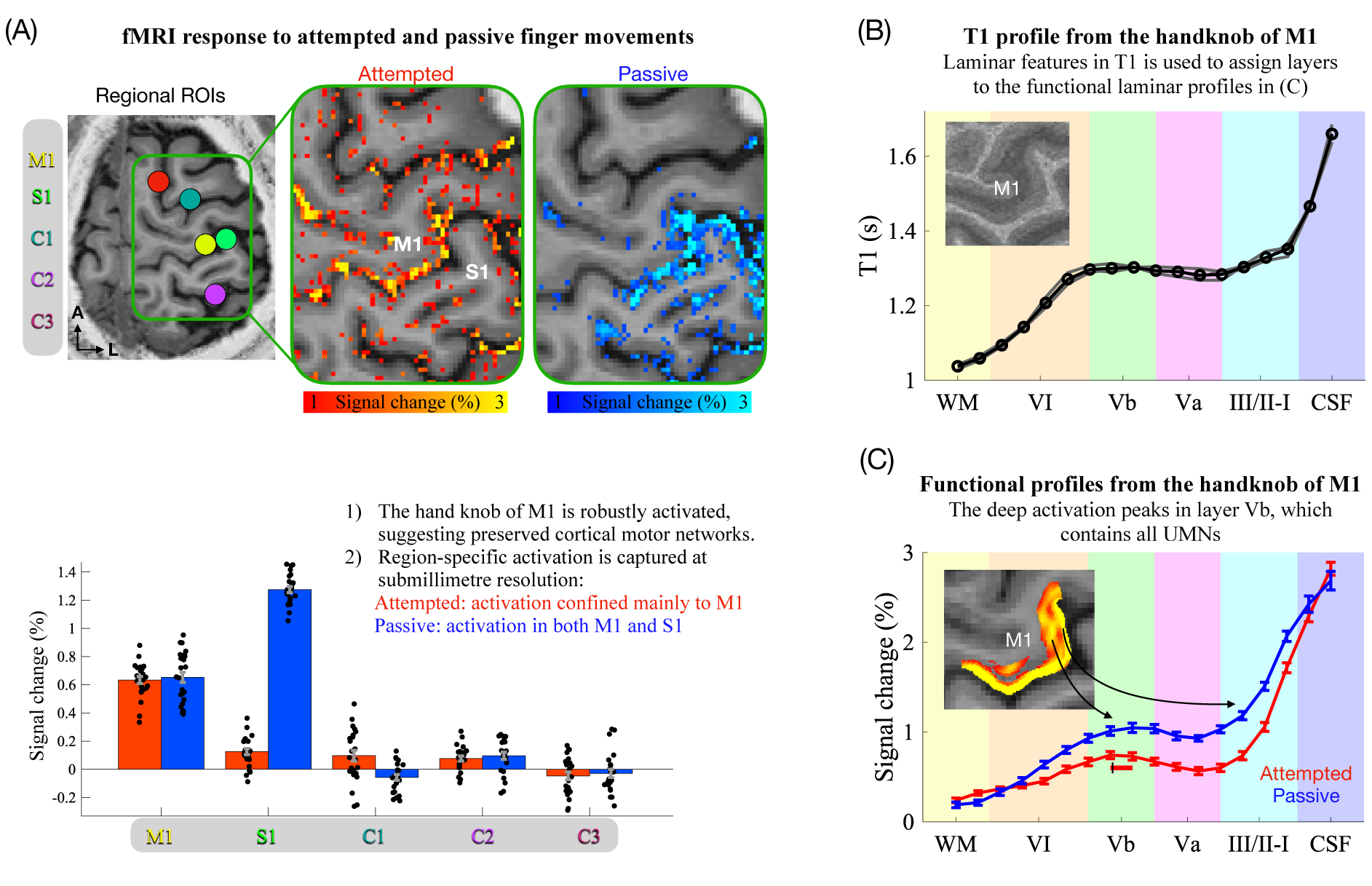
Cortical activation evaluated with submillimetre fMRI. (**A**) BOLD responses to attempted and passive finger movements shown in a representative slice (more slices are shown in Fig. S2A). Activation of each condition was quantified as the average response within ROIs defined in M1 and S1, and compared with three control regions (C1, C2, C3) assumed to be largely inactive for these tasks. ROIs were cylindrical and extended through eight total axial slices. Bar graphs represent the mean and SEM across trials (N=24, represented by black dots). (**B**) Depiction of the quantitative T1 laminar profile obtained from the average of the three MP2RAGE-highres scans (Fig. 1D, Materials and Methods). Based on previous studies^18,31,32^, the depth of layer Va was estimated as the first half of the T1-plateau, the latter half of the plateau was used as a landmark for layer Vb and the sharp decrease in T1 denoted the beginning of layer VI. The beginning and end of the plateau were determined from the first derivative of the T1 profile (Fig. S3). The inset shows the mean T1 image across scans (N=3), thin grey lines represent individual scans, the bold black line is the mean, and dots denote the 18 sampled depths. (**C**) Depiction of the laminar BOLD profiles. Attempted movements were associated with a peak in deep layers, co-localising with the estimated position of layer Vb (the vertical black line and the horizontal red line represent bootstrapped mean and CI95, respectively, for the depth of the deep activation peak, see Materials and Methods). This demonstrates sustained activation in the output layers of M1, possibly driven by preserved UMN activation. The inset shows the activation map for attempted movements following within-layer smoothing highlighting activation stripes in both superficial and deep layers. Error bars reflect SEM across trials (N=24). Note that the 18 bins do not reflect the effective spatial resolution (upsampled grid, see Materials and Methods).

### Laminar analysis reveals preserved layer Vb activation

The functional laminar profiles are depicted in Fig. 3C (Fig. S2B illustrates the profile extraction procedure, Fig S2C depicts profiles from individual trials and runs), showing significant activation in both superficial and deep layers (BOLD-responses were significantly larger than 0 across all grey matter depth-bins during attempted movements as evaluated with permutation testing, see Materials and Methods). The profiles are biased towards the cortical surface due to unwanted draining veins^29,30^. This signal dispersion somewhat challenges interpretation in superficial layers but leaves the deep layer signal largely unaffected^29,30^. Notably, the peak in deep layers co-localised with the estimated position of layer Vb as estimated from MP2RAGE-highres (Fig. 3B).

## Discussion

Using submillimetre resolution fMRI in the locked-in stage of ALS, we found robust M1 activation in response to attempted and passive finger movements, suggesting sustained motor and somatosensory processing, respectively. Furthermore, attempted movements were associated with a deep layer activation peak localised to layer Vb, providing evidence of preserved activation in the output layers of M1 despite years of paralysis. In the following sections, we discuss the potential involvement of UMNs in driving this signal and highlight the capability of laminar fMRI to study in vivo human ALS pathology with unprecedented resolution.

### Layer-dependent M1 activation in a locked-in-stage ALS patient

Passive movement of the index finger robustly activated the hand knob of contralateral M1 (Fig. 3A). This was expected as M1 is known to be an integrate component in proprioceptive networks^33,34^, and proprioceptive function is largely unaffected in ALS patients^1^. Attempted movements elicited comparable levels of activation (Fig. 3A), which is perhaps more surprising given the focal targeting of the motor system in ALS. That being said, one of the rare fMRI investigations in a locked-in-stage ALS patient similarly found increased BOLD-responses in the motor cortex during attempted movements^3^. Furthermore, movement related cortical potentials, requiring somewhat intact motor networks, have been detected with EEG in paralysed ALS patients^4^. Accordingly, the aggressive neurodegeneration of ALS does not necessarily fully extinguish cortical motor networks (but see^23,35^). This is promising for future therapeutic targeting to restore motor function, particularly if it extends to the output neurons of motor cortex. This was explored here using laminar analysis.

Attempted movements activated both superficial and deep layers with a distinct activation peak in the deep end (Fig. 3C), reminiscent of that seen in healthy individuals for actual movements^18–20,36–40^. In line with known input/output organisation of M1 (Fig. 1A) and co-localisation with layer Vb, the deep layer peak is widely believed to mainly reflect corticospinal motor output emerging from UMNs^2,6,18–21,41,42^. This interpretation is compatible with empirical evidence that activation is confined mainly to superficial layers in tasks presumably devoid of corticospinal output, including tactile stimulation (somatosensory input)^18,20^ and motor imagery (motor input)^6,19^. One potential exception is proprioceptive stimulation, for example induced by passive movements, which has been shown to activate deep layers of M1^21^. This was also found here in both the patient (Fig. 3C) and in HC (Fig. S1) and may reflect proprioceptive context delivered from superficial to deep layers to guide impending motor output. Alternatively, it may reflect corticospinal drive being present even during passive movements (discussed in detail in^21^). Regardless, afferent proprioceptive inputs could not have affected the deep layer signal during attempted movements in the patient, as the finger was not overtly moving.

Accordingly, we interpret the observed deep layer signal during attempted movements to predominantly reflect activation in functional UMNs, supported by the spatial congruence between the deep peak and the estimated position of layer Vb (Fig. 3B-C). This is in line with post-mortem research showing that a substantial portion of UMNs can maintain their principal structure at the end-stage of ALS^43–47^. We hypothesise that for some late-stage ALS patients, a considerable fraction of structurally preserved UMNs remain functional despite years of paralysis but fail to generate overt movement due to axonal degeneration or LMN depletion^1^. We cannot rule out, however, that the signal in layer Vb could alternatively have been driven by, e.g., preserved metabolic responses to synaptic input rather than spiking or by non-corticospinal neurons.

### The value of submillimetre fMRI in ALS research

The existing fMRI literature in ALS is largely characterised by use of supramillimetre spatial resolution, spatial smoothing and spatial normalisation. These strategies enhance SNR at the cost of potential signal contamination across neighbouring areas (illustrated in Fig. S4). Disease-related interpretation could thereby be misled as neighbouring regions may have different pathological signatures. In contrast, submillimetre resolution fMRI offers the possibility to resolve specific areas of interest (Fig. 3A), which is critical for capturing spatially localised pathological features. In this vein, Northall and colleagues recently demonstrated the novel insights to be gained on ALS by zooming in to cortical pathology at the mesoscopic level using laminar structural MRI^31^. The present study builds on that by demonstrating the attainability and value of layer-dependent functional recordings in ALS with laminar fMRI. These approaches could ideally be combined to outline the spatiotemporal neurodegenerative proliferation of the disease as advocated by McColgan et al^2^.

### Limitations

Our investigation into UMN preservation in late-stage ALS is based on a single patient. It is noteworthy, however, that our patient was not slowly-progressing (reliant on respiratory assistance 4 years post-diagnosis), had extensive signs of UMN-involvement (Fig. 2B, Materials and Methods) and had suffered from the disease for 6 years - longer than most brains studied post-mortem. These factors favour the notion that a notable proportion of late-stage ALS patients, beyond those primarily exhibiting LMN-dominated pathology, may retain functional UMNs. Nonetheless, validation in more patients is needed to make generalised population claims, especially considering the vast heterogeneity of the disease.

Another limitation, related to laminar acquisitions in ALS, is potential cortical atrophy, iron deposition, vascular deficiency, etc. Although cortical atrophy and pyramidal tract depletion did not clearly manifest in our patient (Fig. 2A,C), the entire cortical ribbon or corticospinal tract might be largely dissipated in severe cases^23,51^. However, this presumably mostly applies in late-stage patients, supported by the observation that the principal laminar architecture appears largely preserved at early stages^31^. Moreover, in instances of severe atrophy where functional recordings might be infeasible, the extent of atrophy itself would often serve as a more direct marker of neurodegeneration and thus reduce the need for such recordings.

## Conclusion

The present study revealed robust activation in the deep output layers of M1 during attempted finger movements in a locked-in stage ALS patient, interpreted to be driven primarily by functionally preserved UMNs. This observation demonstrates a sustained ability to engage motor cortical output layers despite years of paralysis, which fuels hope for future therapeutic targeting. Future measurements from more patients are needed to establish how these findings generalise across disease stages and ALS phenotypes. Finally, our findings highlight an avenue for evaluating mesoscale pathology in vivo using laminar fMRI - an endeavour that has historically been beyond reach but holds promise to shed new light on the perplexing pathology of ALS.

## Materials and Methods

### Participants

One patient (male, age: late 40s) suffering from ALS was included and was diagnosed according to the diagnostic criteria at the time (El Escorial). Exclusion criteria were claustrophobia, other severe medical or neurological diseases unrelated to ALS, implants incompatible with MRI, and sensory deficits in the relevant hand. The patient had suffered from ALS for 6 years, and symptom onset was 7 months prior to diagnosis. Due to severe respiratory failure the patient was initiated on invasive mechanical ventilation 4 years after diagnosis and was at the time of this study in the locked-in stage in which communication was only possible using an eye-tracking communication device or via eye blinking. He was completely paralysed in both hands with a score of 0 in all upper extremity muscles on the Medical Research Council scale, and only able to elicit very weak contractions in lower limbs and in facial muscles. These conditions posed significant challenges for MRI scanning (Fig. 1B), which in combination with the rarity of locked-in stage ALS patients, presumably explain the scarcity of MRI data in this clinical population (but see^3,23^).

At the beginning of the study, a neurological examination was conducted by a consultant neurologist (last author). On prior exams, extensive UMN findings were present: hyperactive biceps-, triceps-, brachioradialis- and achilles-reflexes, bilateral extensive plantar response, positive jaw-jerk reflex, and bilateral foot-clonus. At the time of this study, no UMN findings were present, presumably due to masking by advanced LMN depletion. Also, there were no clinical signs of somatosensory deficits. The patient’s total ALSFRSr score was 1 (Salivation subitem score of 1). He was taking Riluzole (50 mg) twice daily, 20 mg of Citalopram daily, and 0.2 mg subcutaneous Glycopyrronium bromide (for excessive saliva).

A healthy control subject (HC, male, age: mid 50s) was additionally enrolled to provide reference results for the patient data in a healthy individual. Exclusion criteria were claustrophobia, severe medical or neurological disease, alcohol or drug abuse, and implants incompatible with MRI.

Subjects were carefully informed and provided written consent according to the Declaration of Helsinki (a caregiver signed on behalf of the patient after consent was given using an eye-tracking device). The study was approved by the regional ethics committee in Central Denmark Region, Denmark (Study ID 1-10-72-28-23).

### Patient preparation for MRI

The patient was carefully instructed about the imaging procedures, and the patient would respond whether instructions were understood by eye blinks. He was moved from a personal wheelchair to the scanner bed by a hospital porter team, assisted by a lift. The personal artificial ventilation device was replaced with an MR-compatible device, and the patient was hand ventilated by an anaesthesiologist during the transition. The patient’s normal dosage of Glycopyrronium bromide (0.2 mg) was administered to block saliva. As an alternative to the emergency ball, consecutive fast blinks were used to signal if something was wrong. This was monitored throughout the whole imaging session by a neurologist sitting behind the scanner with direct visual access to the patient’s eyes. Physiological parameters (heart rate, carbon oxide and oxygen pressure) were monitored by an anaesthesiologist. Before each imaging sequence, the patient was asked whether everything was okay and if the session should proceed or stop.

### Imaging protocol

MRI images were obtained using a 3T Siemens MAGNETOM Prisma scanner (Siemens Healthineers, Erlangen, Germany) equipped with a standard NOVA head coil with 32 receive channels (NOVA Medical, Wilmington, MA, USA) and a SC72 body gradient coil. The following refers to images acquired in session 1 of the patient (see Fig. 1C for overview). MP2RAGE^52^ was acquired as an anatomical reference with parameters: voxel size=0.9 mm isotropic, matrix size=192×240×256, iPAT=2, Partial Fourier=6/8, TE=2.87ms, TR=5000ms, TI1=700ms, FA1=4°, TI2=2500ms, FA2=5°, echo spacing=7.18 ms. Functional images were acquired with a 3D gradient-echo EPI sequence^53^ and parameters: voxel size=0.82 mm isotropic, TR=2200 ms, TE=27 ms, iPAT=3, Partial Fourier=6/8 (zero-filling reconstruction), FA=45 degrees, and matrix size=176×176×26. Both magnitude and phase images were reconstructed and coil combined using adaptive combine with the prescan normalise setting. We additionally acquired a series of images with SS-SI-VASO^54^ from which an EPI-distortion-matched image with high MP2RAGE-like contrast can be computed to facilitate accurate registration of MP2RAGE to EPI space^18,55,56^. Parameters were: voxel size=0.82 mm isotropic, TR=4739 ms, TE=27 ms, iPAT=3, Partial Fourier=6/8, FA=18 degrees, and matrix size=176×176×26, volumes=100. These three sequences were used as part of the fMRI setup as in our previous study^36^. In the present study, we additionally acquired the following structural images due to their potential sensitivity to ALS pathology^24^: a T2-weighted FLAIR image (voxel size=0.9 mm isotropic, matrix size=256×256×192, TR=5 s, TE=387 ms) and a T2*-weighted gradient-echo image (voxel size=0.72×0.57×1.2 mm^3^ (reconstructed as 0.57×0.57×1.2 mm^3^), matrix size=336×384×80, TR=28 ms, TE=20 ms).

In session 2, we acquired MP2RAGE images with 0.45 mm isotropic resolution used for assigning layers to laminar profiles (MP2RAGE-highres, Fig. 1D), which had parameters: matrix size=480×512×128, no GRAPPA acceleration, Partial Fourier=6/8, TE=3.82 ms, TR=5000 ms, TI1=700 ms, FA1=4°, TI2=2500 ms, FA2=5°, echo spacing=8.96 ms. For HC, we ran the same sequences as in session 1 of the patient.

### Functional MRI experimental design

Two fMRI runs were scanned in the first imaging session of the patient, each with a duration of ∼18 minutes (480 volumes, excluding dummy scans to reach steady state magnetisation). The imaging slab was placed to be approximately perpendicular to the cortical surface of the M1 hand knob in the left hemisphere (Fig. 1C). Each run consisted of blocks of attempted movements, passive movements, and rest. The current condition was indicated to the patient by blinking text saying either “You tap”, “Passive tap”, or “Relax”, presented on an MR-compatible screen placed inside the scanner. Three-second countdowns were implemented to indicate upcoming transitions between blocks. For passive movements, an experimenter was tapping the patient’s right index finger (frequency ∼2.5 Hz) while the patient was asked to fully relax and focus the gaze on the blinking text. These movements were performed by pulling a string attached to the finger. For attempted movements, the patient was asked to attempt to move/tap the right index finger at a frequency and amplitude matched to that of the passive condition (practiced in advance). Resting blocks consisted of relaxation with the gaze focused onto the blinking text. Each block had a duration of 22 seconds and were organised as attempted-rest-passive-rest (representing one trial) which was replicated in 12 trials in each run (Fig. 1C). Head motion was minimised through careful instruction, inflatable padding placed around the head, tape on the forehead for haptic feedback, and wrapping with a heavy blanket around the body. The procedures were identical for HC except attempted movements were replaced by active movements (overt movement).

### Pre-processing of fMRI data

Functional data was pre-processed as in Knudsen et al.^36^. Specifically, reconstructed magnitude and phase images were first denoised using NORDIC (commit 74999d6, downloaded 27-04-2022), which uses principal component analysis to remove components from the data that cannot be distinguished from zero-mean gaussian noise^57,58^. This was done on the complex valued data, without an appended noise volume and with ARG.factor_error=1.15 (default 1), which scales the noise threshold for more aggressive denoising (more components are removed). The purpose was to increase tSNR and thereby the effect of phase regression (see below)^36^.

Motion parameters were estimated from the denoised magnitude timeseries and applied to both magnitude and phase data using SPM12 (Functional Imaging Laboratory, University College London, UK). Estimation was optimised around the hand knob using SPM’s weighting mask option. Phase images were converted to interpolatable real and imaginary components prior to reslicing, and then converted back to phase values before being temporally unwrapped using MATLAB’s (Mathworks Inc.) *unwrap* function.

Phase regression was then applied to reduce the effect of unwanted large draining veins^59^. Briefly, this method relies on the observation that macrovasculature is associated with task-evoked signal modulations in both magnitude and phase images, whereas microvasculature lacks a phase response due to its random vessel orientation. Hence, macrovascular effects can be reduced based on a subtraction of the linear fit between magnitude and phase timeseries from the magnitude timeseries^59^. We recently evaluated NORDIC and phase regression for a largely identical setup^36^. Nevertheless, to rule out that these analysis strategies invalidated our findings, we present the results without NORDIC and phase regression in Fig. S5.

Finally, the MP2RAGE image was registered to EPI space. We first prepared the SS-SI-VASO image which was distortion-matched with respect to the functional acquisitions and had T1-weighted contrast (VASO T1-weighted)^18^. It thus functioned as a template for high-quality registration^55,56^. Preparation included motion correction performed separately on nulled and not-nulled images, which were subsequently aligned to the first image of the functional acquisitions in a single interpolation step using SPM12. These separate timeseries were then concatenated and the signal variability across all timepoints was used to compute the final image. The MP2RAGE image was aligned to the VASO T1-weighted image (and thereby to the functional acquisitions) in the following steps. First, initial parameters were estimated from manual alignment in ITK-snap^60^. These parameters are used as a starting point for subsequent rigid, affine and non-linear (SyN algorithm) registration steps in ANTs^61^. The resulting alignment was carefully assessed by visual inspection to ensure sufficient quality for laminar analysis.

### Activation maps

AFNI^62^ was employed to univariately compute activation maps using the GLM framework^63^. The design matrix consisted of paradigm regressors convolved with a canonical hemodynamic response function. A separate parameter (beta) was estimated for each trial, yielding 24 betas per condition. Beta values were scaled to percent signal change with respect to rest.

### ROI definition

For the evaluation of BOLD-responses in M1, S1, and control regions (Fig. 3A), ROIs were defined anatomically. Specifically, ROIs were selected as cylinders (radius=20 voxels on an upsampled grid with 0.2 mm in-plane resolution) spanning eight slices. For M1, the cylinder was placed on the lateral side of the hand knob, identified from its characteristic omega-shaped landmark^64^. The S1 ROI was placed on the opposite side of the central sulcus, slightly lateral to the M1 ROI^65^. Control regions were defined in random cortical regions outside the sensorimotor areas, expected to show no/minimal activation during attempted/active and passive movements. The cylinders were finally binarized by a grey matter mask obtained with CAT12 (http://www.neuro.uni-jena.de/cat/).

M1 ROIs used for extraction of laminar profiles were defined as follows (illustrated in Fig. S2B). First, borders between white matter/grey matter and grey matter/cerebrospinal fluid were obtained from an initial segmentation of the MP2RAGE image by CAT12, which guided subsequent manual segmentation around the hand knob. Segmentation was performed on an upsampled grid to minimise edge effects (0.2 mm in-plane). These borders were used to compute a depth map and a column map with LAYNII’s *LN2_LAYERS* and *LN2_COLUMNS,* respectively^66^. Timeseries from each of the two runs were then smoothed within columns using *LN2_LAYER_SMOOTH* to elevate SNR, and to avoid laminar information influencing the ROI selection. Note that superficial layers of the column map were discarded to minimise mixing with spatially unspecific pial vein signals. A within-column smoothed activation (beta) map was generated per run as the average of all corresponding betas, reflecting the mean activation across conditions (only used for ROI definition). Two ROIs were then defined, one based on the area showing a strong BOLD-response in the map of the first run, and another based on the map of the second run. This was done to utilise all trials for both localisation and inference without circularity (see “Laminar analysis” below). The search space was limited to the upper and lateral part of the hand knob, namely area BA4a, for comparability with previous laminar M1 studies^18–20,36–40,67^. ROIs had to be contiguous, i.e., without holes, and ended up spanning 6-9 slices (an example slice is shown for the patient in Fig. S2B).

### Laminar analysis

Functional laminar profiles (Fig. 3C) were computed by first dividing the depth map into 18 equidistant layers. Importantly, these layers do not refer to the underlying biological layers, and do not reflect the effective spatial resolution (spatial dependency across layers/bins). At a nominal resolution of 0.82 mm isotropic, and with an approximate M1 cortical thickness of 4 mm in the patient, we expect to sample the thickness with only about 4-5 data points/voxels. Beta values of all ROI-voxels were averaged within each layer for each condition. Importantly, for the ROI defined using trials from the first run, the layer profile was computed from independent trials of the second run, and vice versa for the other ROI. These run-specific profiles were then averaged yielding one laminar profile per condition. All 24 trials per condition could thereby be utilised to both localise the ROI and for statistical inference without circularity^68^.

To highlight layer-dependent features, activation maps were smoothed within layers using LAYNII’s LN2_LAYER_SMOOTH with FWHM=1 mm (only used for visualization purposes in Fig. 3C, and Fig. S2B).

### Generating quantitative T1 profiles to estimate layer positions

Quantitative T1-value maps from the three MP2RAGE-highres images (Fig. 1D) were computed as previously described^52^. Parameters for aligning the second and third run to the first were estimated using ANTs registration (performed on the UNI images). The resulting mean UNI image was used to estimate parameters for registration to the MP2RAGE of the first session (co-registered to EPI space). Using the combined transformations, all three quantitative T1 images were aligned and resliced to EPI space of session 1 in a single interpolation step (upsampled 0.2 mm in-plane grid). Laminar profiles of T1-values were then computed as described for the functional profiles, using the same ROIs.

### Statistical analysis

For all inferential tests, effects were deemed significant at α = 0.05.

Paired two-sided t-tests were used to assess whether BOLD-responses in M1 and S1 were significantly different from the average response across the three control regions. This was done across the 24 trials for each condition. Four total tests were performed in each subject (2 conditions by 2 regions) and corrected for multiple comparisons with the Holm-Bonferroni approach^69^. The mean (±SEM) response of each condition and region is depicted with bar graphs in Fig. 3A (patient) and Fig. S1 (HC). The mean and 95% confidence interval of the most relevant comparisons are reported in the results section. Detailed results are shown in Table S1.

Cluster based permutation tests^70^ were used to assess whether BOLD-responses corresponding to each of the 18 depth-bins (laminar profiles) were significantly different from 0. This strategy was used here, rather than Holm-Bonferroni correction, which might be overly conservative at this number of comparisons. Testing was done across single-trial profiles and separately for each subject and condition. Specifically, in each permutation, the 24 single-trial profiles were randomly multiplied by 1 or -1 as, under the null-hypothesis, each profile could have originated from the contrasts condition>rest and condition<rest with equal probability. P-values were then computed for each bin (across randomly inverted single-trial profiles) using two-sided one-sample t-tests. The sum of t-values in a cluster (defined as range of consecutive bins with uncorrected p-values < 0.05), was then extracted (largest sum if more than one cluster). A null distribution was generated by repeating this process in 10.000 permutations, from which two-sided p-values could be derived for a given cluster.

Layer Vb activation during attempted movements was used here as evidence for potentially preserved UMNs in a late-stage ALS patient. Therefore, confidence limits for the depth of peak activation in deep layers during attempted movements were obtained using bootstrapping^71^. Single trial laminar profiles (N=24) were resampled across 10.000 iterations with replacement. In each iteration, the depth of peak activation in the average resampled profile was computed and 95 % confidence limits were derived from the 2.5 and 97.5 percentiles of the resulting distribution. Since the purpose was to evaluate the spatial certainty of the activation peak in deep layers, the superficial depths (bins 14-18) were discarded in this analysis.

## Data availability

Imaging data associated with this article cannot be made publicly available as anonymisation was not possible given the case-based nature of the study. Data needed to support the conclusions of the study is to the best of our ability made available in the manuscript and supplementary materials. All analysis code will be available at https://github.com/LasseKnudsen1/laminar-fMRI-in-locked-in-stage-ALS.

## Acknowledgements

We would like to sincerely thank the volunteer patient for participating in the study, and the patient care assistant for invaluable help in preparation before scanning. We would further like to thank Ryan Sangill, Dora Grauballe, and Michael Geneser for valuable guidance and assistance during data collection.

## Funding

This project was supported by grants from Dansk Selskab for ALS and Louis-Hansen Fonden. Lasse Knudsen was supported by Sino-Danish Center (SDC). Yan Yang was supported by STI2030-Major Projects 2022ZD0204800 and Beijing Natural Science Foundation (Z210009). Peng Zhang was supported by STI2030-Major Projects (2022ZD0211900, 2021ZD0204200).

## Competing interests

None.

## Notes

### Competing Interest Statement

The authors have declared no competing interest.

