## Supplementary material for "Laminar fMRI in the locked-in stage of amyotrophic lateral sclerosis shows preserved activity in layer Vb of primary motor cortex"

### Supplementary Tables & Figures

**Table S1 – Statistical results corresponding to Fig. 3A and Fig. S1.** CI95: 95% confidence intervals, DOF: degrees of freedom, M1: primary motor cortex, S1: primary somatosensory cortex, HC: healthy control.

| **Fig. 3A – Regional activation patient** | **Mean PSC difference** | **CI95** | **Paired two-sided t-test across trials (N=24)** | **p-value** |
|---|---|---|---|---|
| M1 versus control regions - attempted movements | 0.59 pp. | CI95[0.55;0.63] | t=32.45, DOF=23 | p<0.001 |
| S1 versus control regions – attempted movements | 0.08 pp. | CI95[0.04;0.13] | t=3.59, DOF=23 | p<0.01 |
| M1 versus control regions - passive movements | 0.65 pp. | CI95[0.59;0.71] | t=21.71, DOF=23 | p<0.001 |
| S1 versus control regions – passive movements | 1.27 pp. | CI95[1.23;1.31] | t=60.53, DOF=23 | p<0.001 |
| **Fig. S1 - Regional activation HC** | **Mean PSC difference** | **CI95** | **Paired two-sided t-test across trials (N=24)** | **p-value** |
| M1 versus control regions - attempted movements | 0.50 pp. | CI95[0.35;0.65] | t=7.06, DOF=23 | p<0.001 |
| S1 versus control regions – attempted movements | 0.70 pp. | CI95[0.59;0.82] | t=12.34, DOF=23 | p<0.001 |

| | | | | |
|---|---|---|---|---|
| M1 versus control regions - passive movements | 0.44 pp. | CI95[0.29;0.59] | t=6.26, DOF=23 | p<0.001 |
| S1 versus control regions – passive movements | 0.65 pp. | CI95[0.49;0.81] | t=8.37, DOF=23 | p<0.001 |

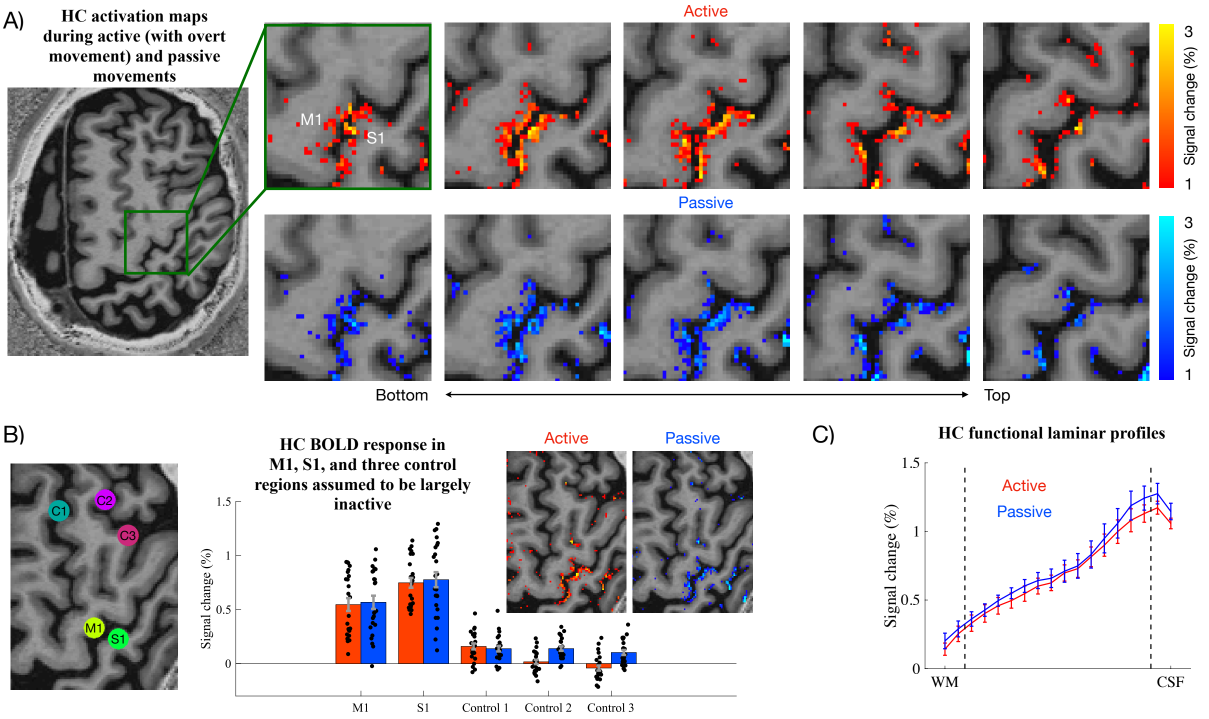


**Figure S1. fMRI results in HC.** (**A**) HC's activation maps across five example axial slices. Note that HC performed actual movements of the right index finger (as opposed to attempted movements in the patient), wherefore this condition is termed "active". Both active and passive movements activated both M1 and S1 as expected (see main text). (**B**) Quantitative evaluation of M1 and S1 responses compared to three control regions (as in Fig. 3A for the patient). Bar graphs represent the mean percent signal change in each ROI, error bars reflect SEM across trials (N=24, represented by dots). (**C**) Laminar profiles of each condition suggesting activation in both superficial and deep layers as in the patient. Notably, there is no clear dip in middle layers, which was observed for the patient (Fig. 3C). Previous similar M1 laminar fMRI studies established that distinct peaks in deep and superficial layers, separated by a dip in the middle, is a robust finding in healthy subjects when non-BOLD sequences, such as cerebral blood volume weighted VASO[1], with higher spatial specificity are employed[2–7]. However, it is not visible in all subjects with BOLD due to its lower effective spatial resolution[2,8,9]. Results from more healthy subjects, scanned with the same setup as here (active finger movements), can be found in our previous paper[8].

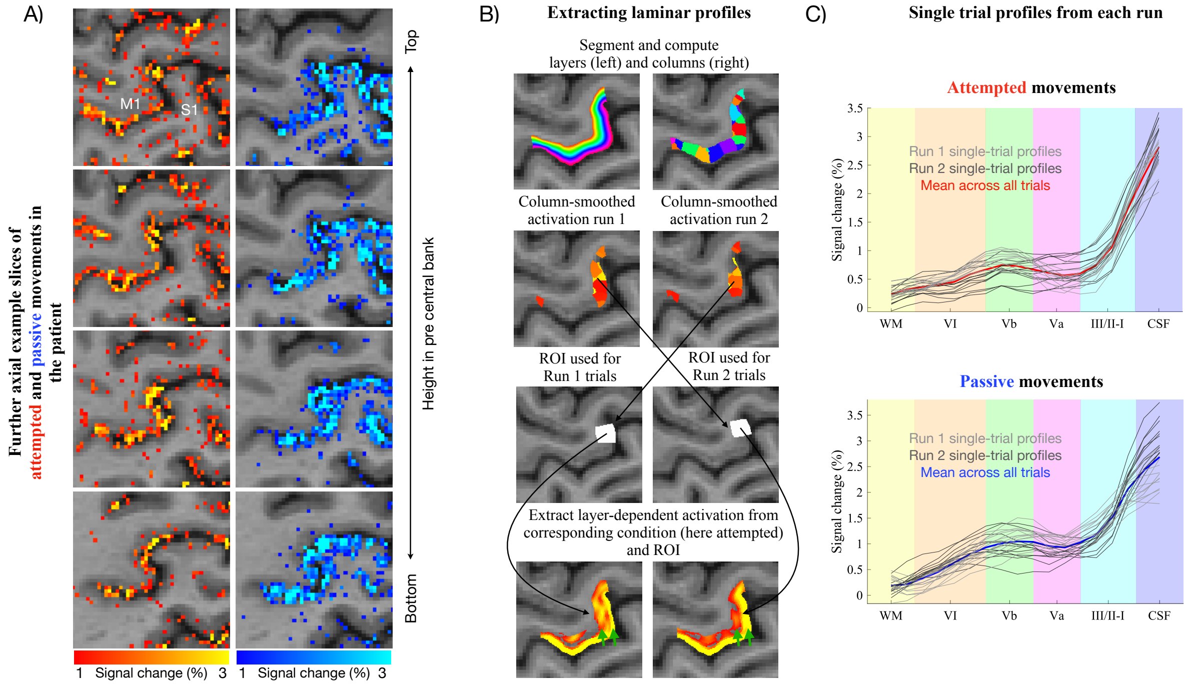


**Figure S2. Activation maps in further example slices, laminar profile extraction and single trial laminar profiles. (A)** The patient's activation maps for attempted and passive movements (as in Fig. 3A), shown in four additional example slices. The expected pattern of activation (mainly M1 for attempted movements, both M1 and S1 for passive movements) was reproducible across axial slices with different heights in the precentral bank of the central sulcus. **(B)** Illustration of ROI definition procedure for the laminar analysis. Briefly, layer and column maps were generated from manually segmented tissue borders around the hand knob of M1. ROI definition was performed by selecting columns that both showed a strong BOLD response (average of attempted and passive trials) and were located within the phylogenetically older and upper part of the M1 hand knob (BA4a) to facilitate comparison with previous laminar fMRI studies in healthy subjects[2–8,10]. To avoid double-dipping, the within-column smoothed activation map of the first run was used to define the ROI for the second run and vice versa. ROIs spanned multiple slices (example slice shown here). Laminar profiles of quantitative T1-values (Fig. 3B) and BOLD-responses (Fig. 3C) were generated by averaging corresponding within-ROI voxel-values in each of 18 equidistant depth-bins/layers. Profiles were extracted separately for each run and then averaged across runs to generate the final laminar profiles. To highlight distinct activation stripes in deep and superficial layers (denoted by green arrows), the functional data was here smoothed within layers. **(C)** Single trial profiles of attempted and passive finger movements in the patient. Light grey and dark grey profiles correspond to single trial profiles from run 1 and run 2, respectively. The coloured profiles correspond to the average across all trials.

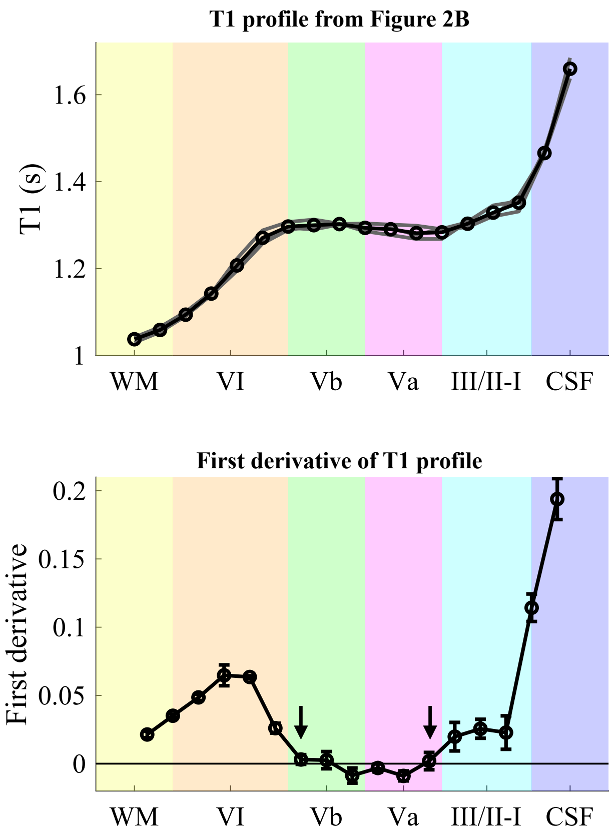


**Figure S3. Using the first derivative of the T1 laminar profile to identify plateau.** The upper panel shows the T1 profile from Fig. 3B. Based on previous studies[2,11,12], the depth of layer Va was estimated as the first half of the T1-plateau, and the latter half of the plateau was used as a landmark for layer Vb. The plateau was demarcated using the first derivative (range of values ~0 between the arrows in the lower panel).

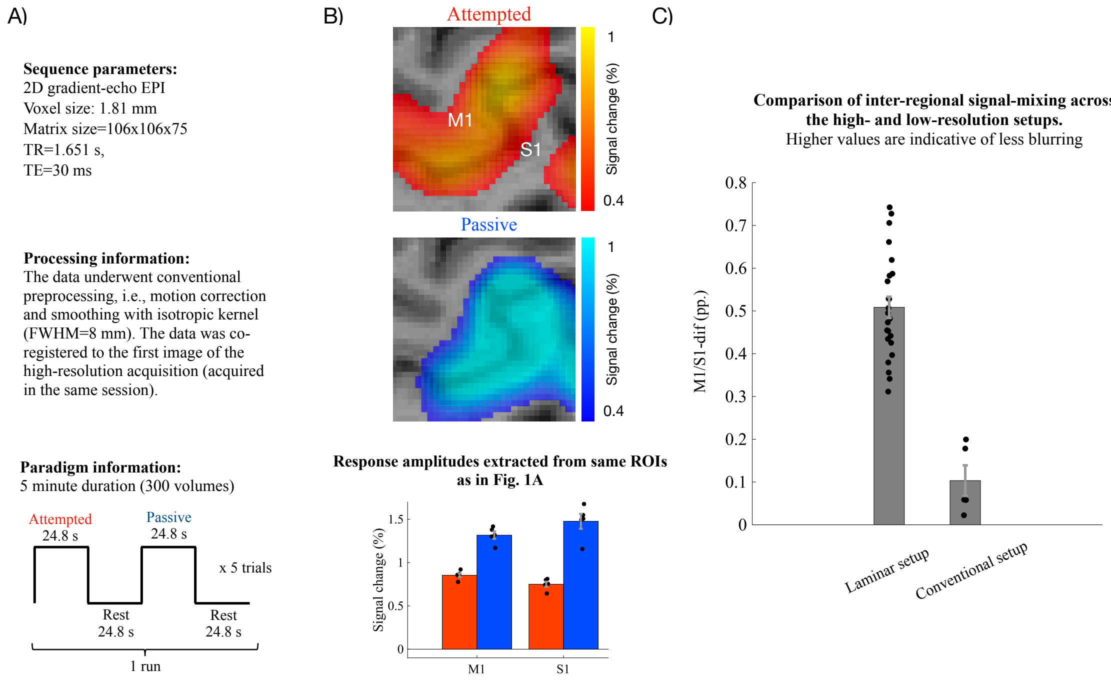


**Figure S4. Illustration of inter-regional signal contamination with conventional fMRI setup.** To illustrate the superior localization power of laminar fMRI setups, an additional fMRI run (outside those depicted in Fig. 1C) was acquired in session 1 of the patient. (**A**) This run was acquired with supramillimetre resolution (1.81 mm isotropic, i.e. >10 times larger voxel-volume than in the laminar setup), and the data was spatially smoothed (FWHM = 8 mm) to represent conventional fMRI setups (i.e. non-laminar), employed in the existing ALS fMRI literature[13,14]. The paradigm was structured as for the laminar setup (Fig. 1C), but with different timings and durations. (**B**) The top panels depict the resulting activation maps. In contrast to the submillimeter resolution data (Fig. 3A), activation maps are here characterized as large blobs, blurring into both M1 and S1. The bar graphs in the lower panel represent the mean and SEM percent signal change across trials (N=5, represented by dots), extracted from the same ROIs as in Fig. 3A. Notably, in contrast to the laminar setup, S1 was robustly activated during attempted movements. (**C**) As argued in the main text, S1 should not be activated during attempted movements due to the lack of somatosensory input without overt movement. Therefore, we quantified the degree of inter-regional blurring as the difference in response amplitude (percent signal change) between M1 and S1 (M1/S1-dif) during attempted movements. M1/S1-dif was on average 5.1 times larger at high resolution (0.51 pp., CI95[0.46;0.56]) than at low resolution (0.10 pp., CI95[0.004;0.20]). This was significant as evaluated with a two-sample, two-sided t-test (mean M1/S1-dif of 0.41 pp., CI95[0.29,0.52], $t(27)=7.11$, $p<0.001$), suggesting that functionally distinct responses between nearby regions were poorly captured at this low effective resolution compared with the laminar setup.

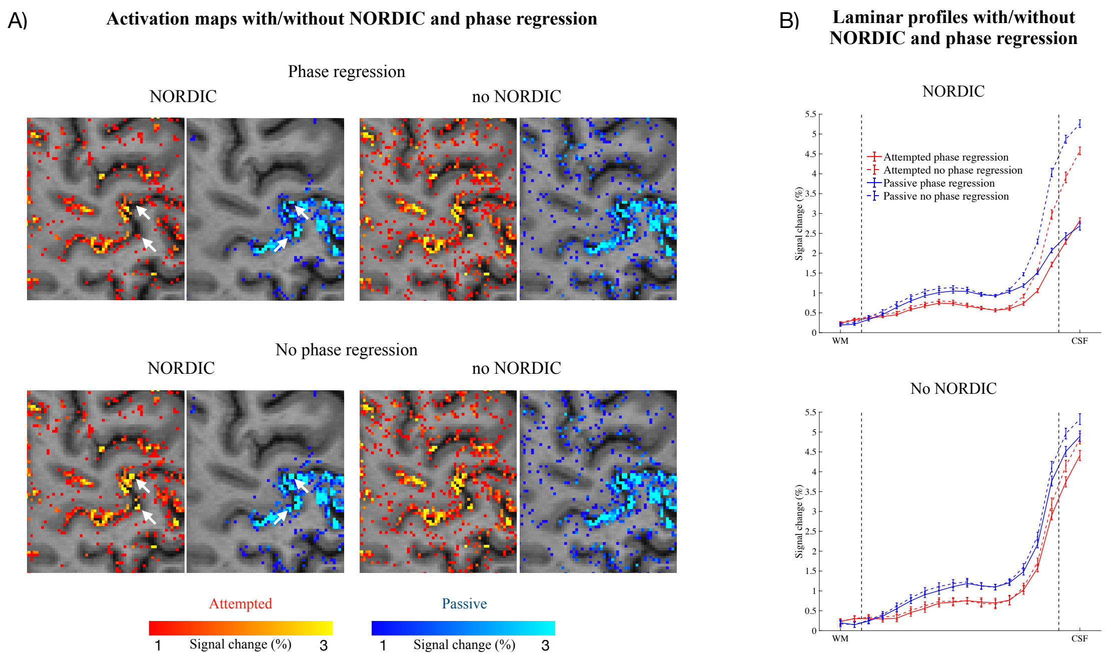


**Figure S5. fMRI results in the patient without NORDIC and phase regression.** (**A**) NORDIC was used to increase the signal-to-noise ratio which is limited at 3T with submillimetre resolution, and phase regression was used to reduce the impact of large veins on obtained BOLD-responses. This setup was validated in our recent study[8]. However, any denoising/filtering method is susceptible to removing signal of interest. To assure these methods did not harmfully affect our data, we compared results across different on/off combinations of NORDIC and phase regression. In this plot, activation maps are shown in an example axial slice for each NORDIC and phase regression combination. NORDIC clearly enhanced signal-to-noise as evident from the reduced false positive activation across the slice. Phase regression suppressed some signal in cerebrospinal fluid (originating from macrovasculature) as highlighted by white arrows, but residual macrovascular influence clearly remained. Importantly, the signal of interest in grey matter of M1 and S1 appears largely identical across all combinations. (**B**) Laminar profiles of all eight combinations. All profiles appear similar, except for the combination of NORDIC and phase regression where the macrovascular bias towards superficial layers is reduced. This is expected since phase regression has limited effects when the signal-to-noise ratio is insufficient (i.e., without NORDIC)[8].

### References supplementary material
